# Understanding proneural–mesenchymal transition using patient-derived glioma stem-like cell (GSC) organoids and engineered extracellular matrix

**DOI:** 10.1101/2025.09.17.676829

**Authors:** Autumn McManis, Charles Ashley Jimenez, Abha Shirolkar, Syed Raza ur Rehman, Sumana Mallick, Malea Murphy, Akhilesh K Gaharwar, Irtisha Singh

## Abstract

Glioblastoma multiforme (GBM) is a highly aggressive, angiogenic WHO grade IV glioma marked by rapid progression, therapeutic resistance, and poor prognosis. A defining feature of GBM is the presence of glioma stem-like cells (GSCs), which reside in specialized perivascular niches and drive tumor progression, recurrence, and therapeutic resistance. The blood-brain barrier, coupled with the complex and dynamic tumor microenvironment, poses significant challenges for both treatment and mechanistic investigation. Current in vitro GBM models inadequately recapitulate the structural and biochemical cues of the native perivascular niche due to the absence of functional vasculature and brain-mimetic extracellular matrix (ECM), limiting their physiological relevance and predictive power. To address the limitations of existing in vitro GBM models, we developed a patient-derived glioma stem cells (GSC) derived Matrigel spheroid system that transitions into organoids and enables integration into engineered microenvironments. Our model incorporates GSC organoids representing proneural and mesenchymal GBM subtypes, a synthetic engineered extracellular matrix (eECM), and endothelial cells (ECs) seeded on the matrix surface. We evaluated the expression of subtype-specific, pro-angiogenic, stemness, and differentiation markers under increasingly complex co-culture conditions. Our results show that Matrigel-derived GSC spheroids progressively differentiate into organoids over two weeks, with significantly enhanced expression of cell-specific markers in the presence of ECs. Encapsulation of these organoids within eECM, combined with EC co-culture, further promoted cellular invasion and induction of GBM associated genes. This *in situ* encapsulation strategy enables real-time observation of GSC behavior in a tunable microenvironment that mimics key features of the native tumor niche. Together, this platform provides a physiologically relevant and modular in vitro system for investigating GBM pathophysiology. It holds promise for uncovering tumor-specific cellular dependencies, studying GSC-vascular interactions, and conducting high-throughput drug screening under controlled, biomimetic conditions.

**Significance:** This study establishes a 3D GSC organoid model with engineered matrix to investigate glioblastoma plasticity and vascular mimicry.

## INTRODUCTION

Glioblastoma multiforme (GBM) is the most aggressive and fatal primary brain malignancy in adults, distinguished by its rapid progression, diffuse infiltration, and resistance to current standard-of-care therapies. Despite over three decades of intensive research and numerous clinical trials, therapeutic advances have yielded only modest improvements, with median survival increasing from approximately 12 to 15 months and a five-year survival rate remaining below 5%^1^. The failure to significantly improve patient outcomes underscores the urgent need for innovative approaches to study GBM pathophysiology and develop effective therapies. Efforts have been made to develop techniques to better image^2^ and monitor physiological changes^3^ happening in a patient’s diseased brain to potentially aid the direction of treatment needed, however there is still much to elucidate regarding the mechanisms driving the development and recurrence of GBM. Current preclinical models are insufficient to recapitulate the complexity of the human disease. In vitro cell-autonomous models lack physiological relevance, while in vivo models are costly, time-consuming, and offer limited experimental tractability. Existing three-dimensional (3D) platforms such as tumor organoids^4^, tumor-on-a-chip^5, 6^, patient-derived xenografts^7^, and genetic mouse models^8^, address certain aspects of GBM biology, but fall short in capturing the broader spectrum of tumor heterogeneity and microenvironmental cues. A clinically relevant, scalable, and tunable in vitro model that better mimics the native tumor niche is essential for advancing both mechanistic understanding and therapeutic discovery in GBM.

GBM is characterized by extensive vascularization and pronounced cellular hierarchies, with glioma stem-like cells (GSCs) occupying the apex. These GSCs are highly tumorigenic, therapy-resistant, and play critical roles in sustaining tumor growth, promoting angiogenesis, and facilitating invasion into healthy brain tissue^9-11^. Within the tumor microenvironment, GSCs reside in specialized niches where they receive maintenance cues from surrounding cells and extracellular matrix (ECM) components^12, 13^. A central component of this niche is the tumor vasculature, composed of brain endothelial cells (ECs) that supply both structural and biochemical support including Notch ligands, ECM proteins, and nitric oxide to regulate GSC behavior^14-16^. Additional perivascular cells include pericytes that reside directly outside the ECs within the basement membrane and astrocytes that connect to the exterior of the basement membrane, both of which play a crucial role in the function of the blood-brain barrier providing the capillary function to pass nutrients and signaling between the vasculature and surrounding cells^17^. Studies suggest that GSCs can differentiate into such vascular supporting cells^12, 18-20^. Direct and indirect interactions between GSCs and ECs within the perivascular niche govern key aspects of tumor progression, including stemness, migration, therapeutic resistance, and cellular kinetics^21-23^. Understanding the regulatory mechanisms within this niche is essential for identifying vulnerabilities that can be targeted therapeutically.

Conventional 2D culture models are unable to fully capture the architectural and biochemical complexity of the GBM microenvironment, often resulting in inconclusive experimental outcomes^24, 25^. In contrast, 3D models such as neurospheres^20, 26^, organotypic cultures^26-29^, and hydrogel-based encapsulation^26, 28^ more closely mimic in vivo disease architecture and cell-matrix interactions. However, most current 3D systems still lack critical components of the native tumor niche such as brain-specific ECM components^30, 31^, perivascular structure^32-34^, and interaction with non-tumor cells^35, 36^, which are essential for maintaining GSC function and therapeutic resistance. Some models have tried to overcome these challenges by incorporating different neuronal and non-neuronal cells within a 3D matrix to recapitulate GBM microenvironment, but there is still significant room to develop 3D models that mimic native biophysical and biochemical heterogeneity^23, 37, 38^. Thus, there remains need for developing next-generation 3D models that incorporate microenvironmental features along with cellular heterogeneity mimicking native GBM while allowing high-resolution analysis and experimental flexibility.

In this study, we present a bioengineered GBM model that integrates patient-derived GSCs into a Matrigel-based spheroid system capable of transitioning into organoids and embedding within tunable, engineered extracellular matrices. This model is designed to facilitate controlled co-culture with endothelial cells along with the molecular interactions. Our prior epigenomic analysis of a panel of 30 patient-derived GSC lines revealed two major subtypes distinguished by enhancer profiles and transcriptional programs^39^. Functional validation through knockdown studies demonstrated subtype-specific effects on tumor growth in vitro and in vivo. These findings suggest fundamental differences in cell identity between subtypes, pointing to distinct therapeutic vulnerabilities. To capture this heterogeneity, we will use representative GSCs from both molecular subtypes to construct patient-specific, engineered GBM models. By integrating subtype-specific GSCs, engineered ECM, and vascular cues, this model provides a tangible platform to study GBM pathophysiology, identify context-specific tumor dependencies, and enable high-throughput drug screening in a physiologically relevant environment.

## RESULTS

### Establishing and characterizing GSC Matrigel spheroid system

To establish the GSC spheroid system, it was essential to characterize gene expression patterns associated with specific subtypes and pro-angiogenic activity. Quantitative real-time polymerase chain reaction (qRT-PCR) was performed to assess the expression of proneural markers (OLIG2, BCAN) and mesenchymal markers (STAT3, RUNX2)^39^, as well as pro-angiogenic markers (VEGFA, ANG1, FGF2). GSCs were first cultured as monolayers and then used to generate spheroids (10,000 cells/spheroid) and Matrigel spheroids (10,000 cells/spheroid with Matrigel; Materials and Methods). Both spheroids and Matrigel spheroids were cultured for 3 or 14 days prior to RNA isolation for qRT-PCR analysis (**Figure 1A**).

**Figure 1.**
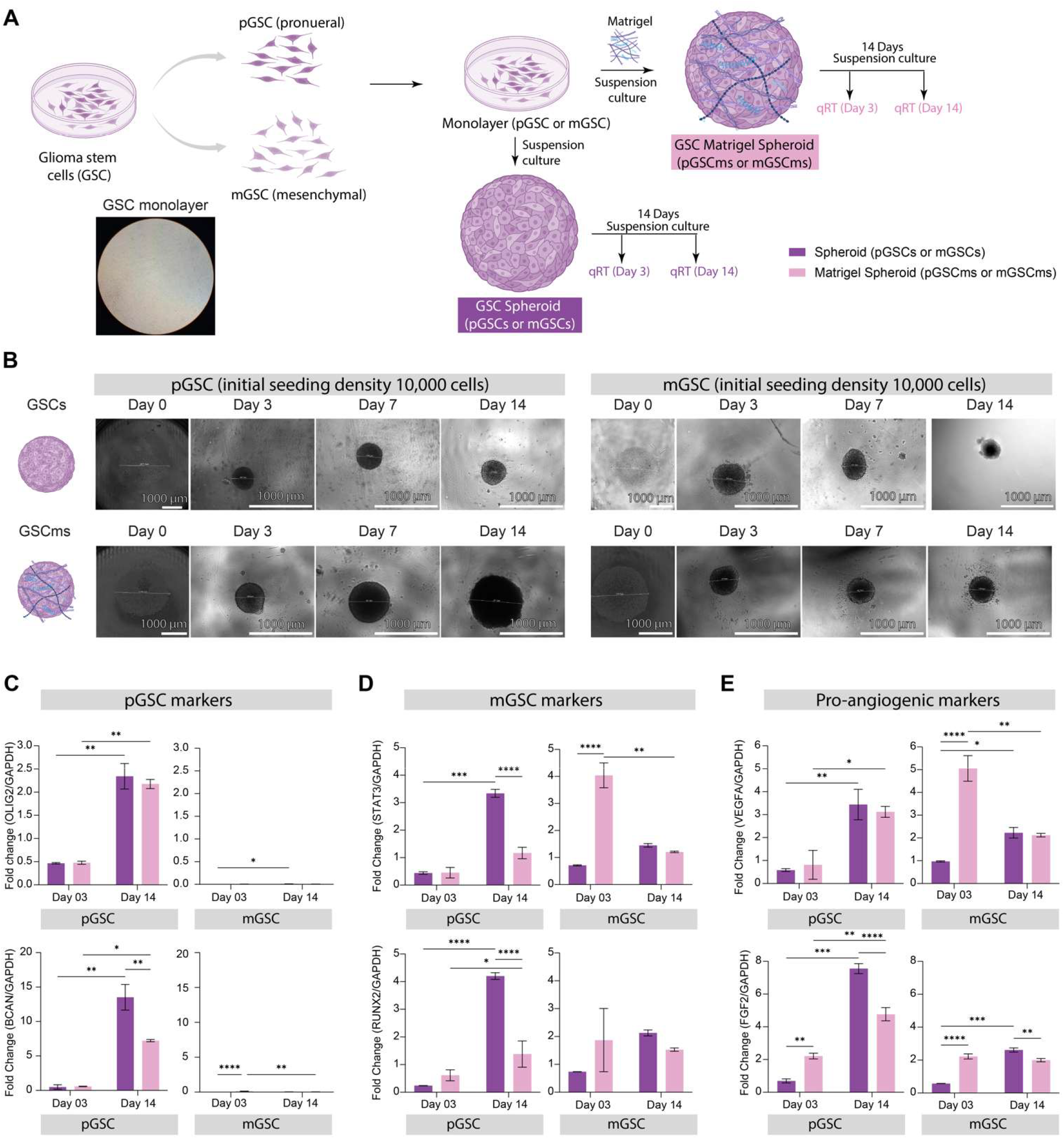
Establishing and Characterizing the GSC Organoid System. (A) Schematic overview of the experimental workflow. Patient-derived glioma stem cells (GSCs) representing either the proneural (pGSC) or mesenchymal (mGSC) subtype were cultured under two conditions: spheroid (pGSCs/mGSCs) and Matrigel spheroid (pGSCms/mGSCms). Cells were harvested after 3 or 14 days for RNA extraction and gene expression analysis. (B) Phase contrast images showing the progression of spheroid and Matrigel spheroid formation over time. Samples were imaged on days 0, 3, 7, and 14. (C–E) RT-qPCR analysis showing fold change in expression of group-specific and pro-angiogenic markers, normalized to GAPDH. (C) Expression of pGSC-specific markers OLIG2 and BCAN, relative to pGSC spheroid day 3 controls. (D) Expression of mGSC-specific markers STAT3 and RUNX2, relative to mGSC spheroid day 3 controls. (E) Expression of pro-angiogenic markers VEGFA and FGF2, relative to its own GSC spheroid day 3 controls. Data are presented as mean ± SD (N = 3). Statistical analysis was performed using two-way ANOVA with Tukey’s post hoc test (*p ≤ 0.05, **p ≤ 0.005, ***p ≤ 0.0005, ****p ≤ 0.00005).

To evaluate the maintenance of a proneural gene expression profile, *OLIG2* and *BCAN* transcript levels were measured in pGSC (proneural Glioma Stem Cells) and mGSC (mesenchymal Glioma Stem Cells) cultured as spheroids or Matrigel spheroids at days 3 and 14 (**Figure 1B**). Oligodendrocyte transcription factor 2 (OLIG2) is a basic helix-loop-helix transcription factor that functions as an oligodendrocyte master regulator^39, 40^. Proneural GSCs express more *OLIG2* than mesenchymal GSCs as evident from GBM, however some studies indicate that mesenchymal GSCs can express OLIG2 and become more migratory^24, 41-43^. Silencing of *OLIG2* in patient-derived proneural GSC appears to result in either proneural-to-classical or proneural-to-mesenchymal transition in different cell lines^44^. Brevican (BCAN) is a protein found in the central nervous system ECM^45^. BCAN is upregulated in GBM, particularly in proneural GBM^45^. In pGSC, both spheroid and Matrigel spheroid cultures showed a significant increase in *OLIG2* and *BCAN* expressions over time (**Figure 1C**). However, when mGSC were cultured as spheroids or Matrigel spheroids, a marked reduction in OLIG2 and BCAN expression was observed over time in Matrigel spheroids. As *OLIG2* and *BCAN* are proneural markers, these findings suggest that Matrigel spheroid cultures are more suitable for maintaining appropriate long-term culture conditions for GSCs.

We next examined the mesenchymal gene expression profile of GSCs cultured as spheroids or Matrigel spheroids by quantifying *STAT3* and *RUNX2* transcript levels at days 3 and 14 (**Figure 1D**). Signal transducer and activator of transcription 3 (STAT3) encodes transcription factors in response to cytokines and other stimuli^46^. Mesenchymal GSC can be identified by their increased *STAT3* expression versus proneural GSC, indicating increased cell motility and malignancy^39^. Runt-related transcription factor 2 (RUNX2) is a transcription factor protein that regulates mesenchymal stem cells towards osteoblast differentiation and skeletal development (bone, cartilage, etc.)^47, 48^. Mesenchymal GSC have increased expression of transcription factors such as RUNX2, poorer patient prognosis, increased affinity towards hypoxic areas, increased resistance to radiotherapy, and more migratory capabilities^39, 49, 50^. In pGSC, *STAT3* and *RUNX2* expression was minimal on day 3 across all culture conditions. In contrast, both spheroids and Matrigel spheroids exhibited a significant increase in *STAT3* and *RUNX2* over time. mGSC showed higher baseline expression of both markers in spheroids and Matrigel spheroids at day 3, with these elevated levels persisting through day 14. Overall, these results indicate that both pGSC and mGSC display increased mesenchymal gene expression under 3D culture conditions^39^. Notably, mGSC Matrigel spheroids exhibited higher mesenchymal marker expression than pGSC Matrigel spheroids, a difference not observed in the spheroid-only condition.

To assess the effect of culture conditions on pro-angiogenic signaling, *VEGFA* and *FGF2* transcript levels were measured in pGSC and mGSC cultured as spheroids or Matrigel spheroids at two timepoints (**Figure 1E**). VEGFA is likely the primary pro-angiogenic factor produced by both proneural and mesenchymal GSC^51^. Proneural GSC with inhibited *VEGF* are more likely to take on more vascular phenotypes by differentiating into pericytes, endothelial cells, or smooth muscle cells^50^. The STAT3/NF-κB pathway is activated by FGF2 (fibroblast growth factor 2, basic FGF, bFGF) to promote angiogenesis^52^. In 3D cultures, both cell types showed a significant increase in *VEGFA* and *FGF2* expression, particularly at day 14. These findings indicate that 3D culture enhances pro-angiogenic gene expression in both GSC subtypes, with Matrigel spheroids providing the most consistent environment for upregulating these factors compared with spheroids. Overall, our results show that Matrigel spheroids are the most effective condition for maintaining and enhancing these profiles compared with spheroids. These optimized conditions provide the foundation for subsequent studies of GSC stemness, proliferation, and hypoxia in the presence of endothelial cells.

### Endothelial Co-Culture Enhances Stemness, Proliferation, and Hypoxia in GSC Matrigel Spheroids

To evaluate the overall health and functional state of GSCs in presence of endothelial cells, immunofluorescence (IF) staining was performed using established markers of stemness (CD133), proliferation (Ki67), and hypoxia (CAIX; Materials and Methods). First, GSCs were expanded as monolayers and subsequently used to generate spheroids (1,000 cells/spheroid) or Matrigel spheroids (1,000 cells/spheroid with Matrigel; Materials and Methods). Both spheroids and Matrigel spheroids were cultured for either 3 or 14 days prior to fixation and IF staining. In addition, Day 14 Matrigel spheroids were employed to establish Transwell® co-cultures with human umbilical vein endothelial cells (HUVECs) and maintained for an additional 3 or 14 days before fixation and IF staining (**Figure 2A;** Materials and Methods). IF staining for each marker was normalized to the nuclear signal (DAPI), enabling direct comparison of marker expression across different spheroid conditions (Materials and Methods).

**Figure 2.**
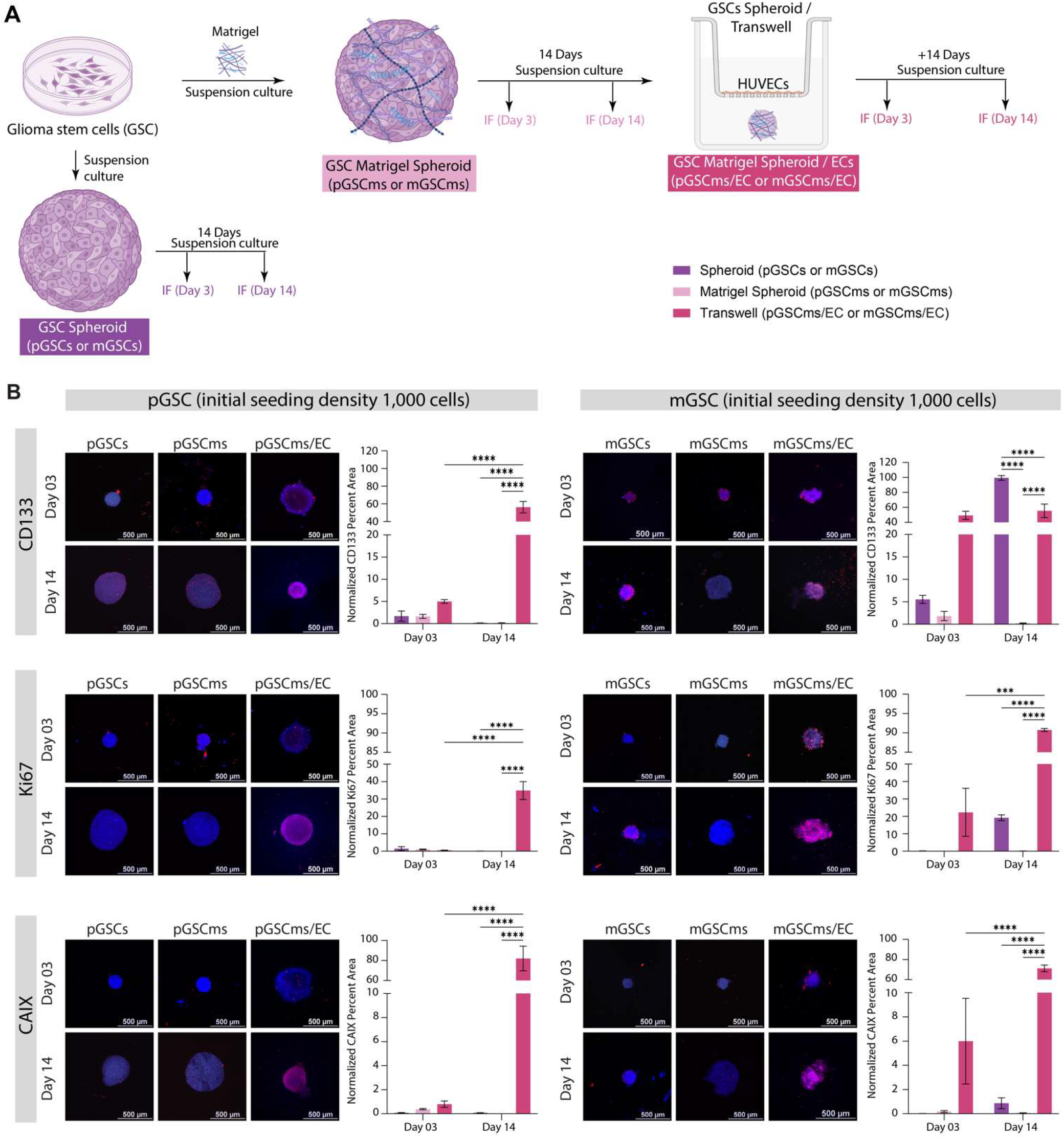
Establishing stemness, proliferation, and hypoxia profiles in GSC organoids co-cultured with endothelial cells. (A) Schematic of the experimental design. Glioma stem cells (pGSC: proneural, mGSC: mesenchymal) were cultured for either 3 or 14 days under three conditions: spheroid (pGSCs/mGSCs), Matrigel spheroid (pGSCms/mGSCms), and Matrigel spheroid co-cultured with endothelial cells via Transwell® inserts (pGSCms/EC, mGSCms/EC). (B) Representative immunofluorescence images and quantification of percent area of fluorescent signal (normalized to DAPI area) for markers of stemness (CD133), proliferation (Ki67), and hypoxia (CAIX) at Days 3 and 14. Marker signal is shown in red; nuclei are counterstained with DAPI (blue). Scale bars = 500 µm. Data are presented as mean ± SD (N = 3). Statistical analysis was performed using two-way ANOVA with Tukey’s post hoc test (*p ≤ 0.05, **p ≤ 0.005, ***p ≤ 0.0005, ****p ≤ 0.00005).

To assess GSC stemness, CD133 expression was first quantified and normalized to nuclear staining (**Figure 2B**). CD133, or prominin-1, is a stem cell surface marker commonly used to identify and isolate GSC^50^. Glioblastoma patients classified as proneural subtype in The Cancer Genome Atlas (TCGA) express higher levels of *PROM1* compared to mesenchymal subtype (**Figure S2A**)^53^. However, there are no marked differences in *PROM1* expression between the glioblastoma patients classified as proneural and mesenchymal subtypes in the Chinese Glioma Genome Atlas (CGGA) ^53, 54^. Both pGSC and mGSC showed baseline CD133 expression at days 3 in spheroid and Matrigel spheroid cultures, with mGSC (mesenchymal) exhibiting slightly higher levels than pGSC (proneural). On day 14, a significant increase in CD133 expression was observed in mGSC spheroid. Notably, co-culture of Matrigel spheroids with endothelial cells (ECs) resulted in a significant increase in CD133 expression for both GSC 1 and mGSC at days 3 and 14, indicating enhanced stemness in the presence of endothelial cells. The increase in stemness of GSCs upon interaction with ECs is consistent with previous findings, which demonstrated that EC interactions promote stemness in vitro^55^.

To evaluate the proliferative capacity of GSCs under different culture conditions, Ki67 expression was assessed on day 3 and day 14. High Ki-67 proliferation index correlates to a highly aggressive glioblastoma, postoperative recurrence, and mortality in IDH-wild-type GBM patients^56^. Both pGSC and mGSC exhibited minimal Ki67 expressions on day 3 across all conditions. By day 14, a significant increase in Ki67 expression was observed specifically in the co-culture condition for both GSC subtypes, with mGSC displaying markedly higher levels of proliferation compared to pGSC. Quantification of normalized Ki67-positive area confirmed that co-culture with ECs significantly enhances proliferation relative to both spheroids and Matrigel spheroids cultured alone. These findings replicate the trends seen in previous studies indicating that endothelial interactions within the Matrigel matrix promote robust GSC proliferation, particularly in the mesenchymal subtype mGSC^57^.

Hypoxia was assessed by CAIX expression across culture conditions. Carbonic anhydrase IX (CAIX) is an acid-base regulatory protein shown to have an increased response in GBM and correlates with an unideal response to anti-cancer therapeutics, such as radiation and chemotherapy, and thus low patient survival rates^41^. Both proneural and mesenchymal GSC express CAIX more than healthy brain tissue; however, mesenchymal GSC have the highest expression, particularly those closest to the necrotic core of a tumor^41, 58^. On day 3, all conditions showed minimal CAIX expression for both pGSC and mGSC. By day 14, CAIX levels remained low in both spheroid and Matrigel spheroid cultures. However, a dramatic increase in CAIX expression was observed in the co-culture condition for both pGSC and mGSC, indicating enhanced hypoxic response due to endothelial interaction. Quantitative analysis revealed a significant elevation in CAIX-positive area in the co-culture group compared to both spheroid and Matrigel spheroid conditions. Altogether, our results reflect previously observed patterns of CAIX upregulation in GSCs, especially mesenchymal GSCs, when in the presence of ECs^41, 58^.

Our results underscore the importance of incorporating endothelial cells into 3D culture models to better recapitulate the glioblastoma microenvironment. The co-culture of GSC Matrigel spheroids with endothelial cells significantly enhances key tumor-associated phenotypes including stemness, proliferation, and hypoxic signaling across both pro-neural and mesenchymal GSC subtypes. This vascularized organoid system provides a physiologically relevant platform for studying GSC biology and offers a valuable tool for evaluating therapeutic responses in a context that more accurately reflects in vivo tumor architecture and signaling dynamics.

### Co-Culture with Endothelial Cells Promotes Lineage Commitment in GSC Spheroids

We next assessed GSC differentiation across varying microenvironments by performing immunofluorescence (IF) staining for lineage-specific markers: GFAP (Glial Fibrillary Acidic Protein, astrocytic marker), MOG (Myelin Oligodendrocyte Glycoprotein, oligodendrocytic marker), CD248 (Endosialin, pericytic marker), and CD31 (Platelet Endothelial Cell Adhesion Molecule-1, endothelial marker). GSCs were first cultured as monolayers and then used to generate spheroids (1,000 cells/spheroid) or Matrigel spheroids (1,000 cells/spheroid embedded with Matrigel). These were maintained for 3 or 14 days prior to fixation and IF staining. Additionally, day 14 Matrigel spheroids were used to initiate Transwell® co-cultures with HUVECs and cultured for another 3 or 14 days before fixation and analysis (**Figure 3**).

**Figure 3.**
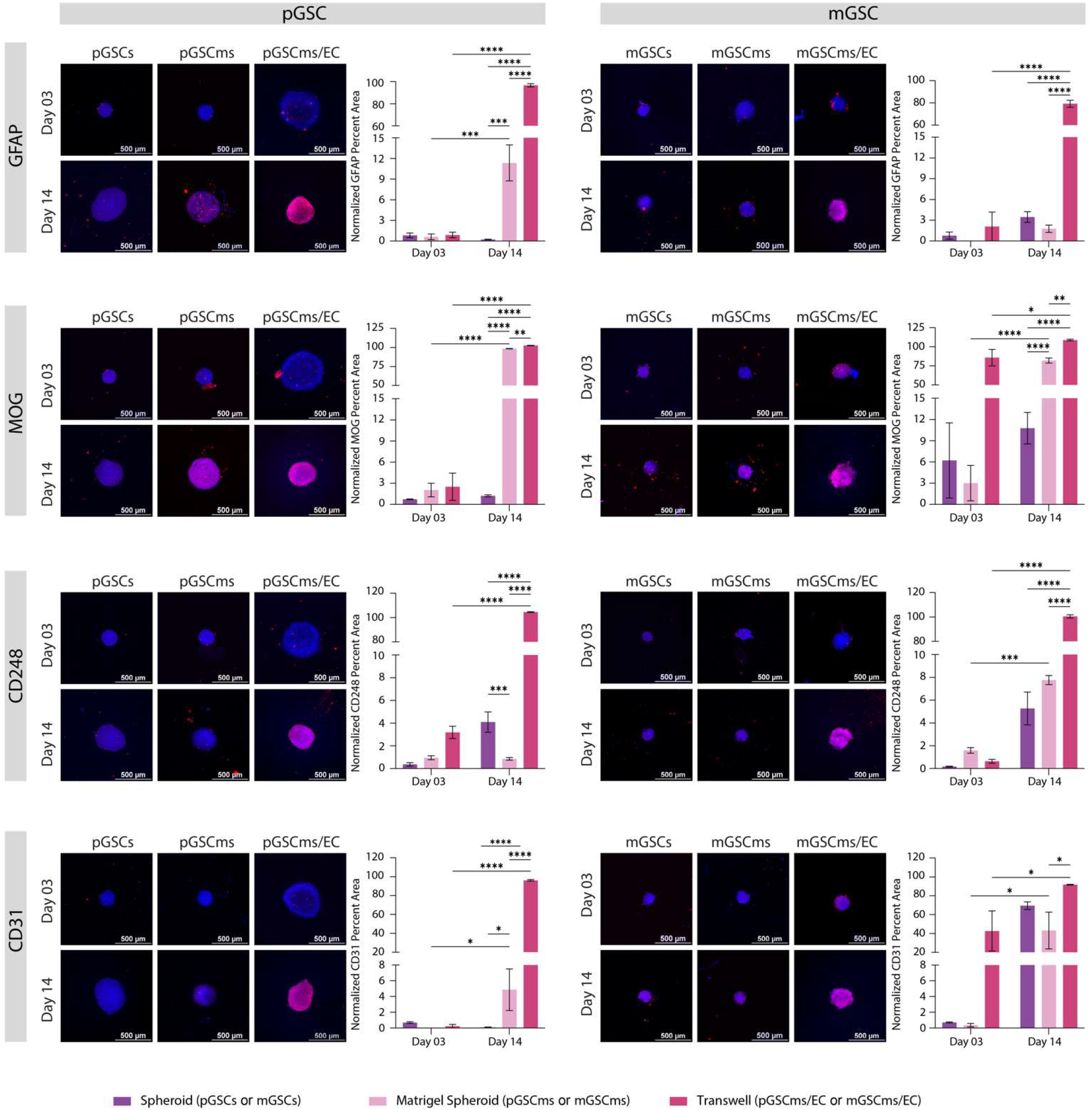
Endothelial co-culture enhances lineage differentiation of GSC Matrigel spheroids. Glioma stem cells (pGSC: proneural; mGSC: mesenchymal) were cultured for either 3 or 14 days as spheroids (pGSCs/mGSCs), Matrigel spheroids (pGSCms/mGSCms), or Matrigel spheroids co-cultured with endothelial cells via Transwell® inserts (pGSCms/EC, mGSCms/EC). Samples were fixed and immunostained for astrocytic (GFAP), oligodendrocytic (MOG), pericyte (CD248), and endothelial (CD31) markers. Representative immunofluorescence images show red signal for the respective markers and blue for DAPI nuclear staining. Quantification of percent area of fluorescent signal (normalized to DAPI area) demonstrated a significant increase in marker expression over time and in the presence of endothelial cells, indicating enhanced lineage differentiation within the Matrigel co-culture system. Scale bars = 500 µm. Data are presented as mean ± SD (N = 3). Statistical analysis was performed using two-way ANOVA with Tukey’s post hoc test (*p ≤ 0.05, **p ≤ 0.005, ***p ≤ 0.0005, ****p ≤ 0.00005).

To evaluate astrocytic lineage commitment of GSCs, we performed IF staining for GFAP across a range of culture conditions. Glial fibrillary acidic protein (GFAP) is an intermediate filament protein, an essential component of the astrocytic cytoskeleton, and is likely linked to GBM progression^57^. Mesenchymal GSC with high STAT3 expression could also differentiate toward astrocytes expressing GFAP^39, 59^. Differentiated proneural GSC increase their GFAP expression and become more migratory^20^. Both pGSC and mGSC showed minimal GFAP expressions at day 3 and 14 in spheroid cultures, indicating limited astrocytic differentiation in the absence of matrigel or endothelial cues. In contrast, Matrigel spheroids exhibited a modest but consistent increase in GFAP expression in pGSC, suggesting that the Matrigel alone supports early astrocytic differentiation in pGSC. Notably, GFAP expression was significantly elevated in Matrigel spheroids co-cultured with HUVECs, particularly on day 14, in both pGSC and mGSC. The results indicate that endothelial-derived cues critically contribute to promoting astrocytic differentiation within the GSC microenvironment. These findings coordinate with previous studies that suggest endothelial-derived signals play a key role in enhancing astrocytic differentiation within the GSC microenvironment^60^.

To assess oligodendrocytic lineage commitment, IF staining for MOG was performed across different GSC culture conditions. Myelin oligodendrocyte glycoprotein (MOG) is a component of the central nervous system and indicates mature, differentiated oligodendrocytes^61^. Although proneural GBMs express more *MOG* than mesenchymal GBMs as evident from TCGA and CGGA patients dataset, MOG is not expressed in pGSC or mGSC (**Figure S2E, Table S1**)^39, 41, 53, 54^. pGSC monolayers and spheroids showed negligible MOG expression at both day 3 and day 14, indicating that simple 3D culture without Matrigel is insufficient to induce oligodendrocytic differentiation. Similarly, mGSC spheroids did not exhibit a significant increase in MOG expression from day 3 to day 14. In contrast, Matrigel spheroids demonstrated a significant time-dependent increase in MOG expression, particularly on day 14, suggesting that Matrigel-derived extracellular matrix cues support early oligodendrocytic differentiation. Notably, MOG expression was further elevated in Matrigel spheroids co-cultured with HUVECs, especially after 14 days of co-culture. This enhancement was observed in both pGSC and mGSC systems. These findings suggest that endothelial cells provide additional instructive signals possibly via paracrine factors or extracellular matrix remodeling—that promote oligodendrocytic lineage specification. Although GBM studies of MOG expression in the perivascular niche are limited, it has been reported that the presence of ECs leads to significantly upregulated OLIG2 which can then lead to increased expression of MOG^55, 62^. The increased MOG expression in co-culture conditions underscores the importance of vascular components in driving maturation toward oligodendrocyte-like phenotypes in GSCs.

We next assessed the pericytic differentiation potential of GSCs by analyzing expression of CD248, a surface marker associated with perivascular stromal cells. Tumor endothelial marker-1 (TEM-1 or CD248) is a transmembrane protein found in pericytes of certain tumors including sarcoma and neuroblastoma and can be expressed on tumor stroma or tumor cells themselves^63^. CD248 plays an important role in tumor growth and blood vessel formation, and a higher expression of CD248 appears to correlate with greater tumor aggressiveness and worse prognosis, making it an attractive target for cancer treatment and imaging^63^. CD248 expression remained low in both spheroid and Matrigel spheroid conditions at early timepoints (day 3). However, a significant increase was observed on day 14, particularly in Matrigel mGSC spheroids. Notably, the most pronounced upregulation of CD248 occurred in Matrigel spheroids co-cultured with HUVECs, where both pGSC and mGSC displayed significantly elevated expression levels on day 14. This suggests that prolonged exposure to endothelial-derived signals, in the context of an extracellular matrix, promotes pericytic lineage commitment in GSCs^12^. The findings are consistent with the hypothesis that endothelial-tumor cell interactions within the perivascular niche can drive mesenchymal transitions and support vascular mimicry^12^.

To assess vascular mimicry and endothelial integration in GSC cultures, we performed immunofluorescence staining for CD31, a well-established endothelial marker. Platelet Endothelial Cell Adhesion Molecule-1 (PECAM-1 or CD31) is a cell surface marker for EC that is typically found at cell-to-cell junctions and indicates blood vessel formation and proliferation^64^. Although it is not yet clear if GSC can differentiate to become EC, some reports indicate that GSC expressing GFAP might differentiate into EC that also express CD31^18^. As expected, no CD31 expression was detected in pGSC spheroid or Matrigel spheroid cultures without HUVECs, confirming the absence of spontaneous endothelial differentiation in proneural-type GSCs. In contrast, mGSC spheroids and Matrigel spheroids exhibited a modest but significant increase in CD31 expression by day 14, indicating a limited capacity for endothelial-like transition in mesenchymal-type GSCs. Strikingly, upon co-culture with HUVECs, both pGSC and mGSC Matrigel spheroids showed robust CD31 expression, which significantly increased from day 3 to day 14. Notably, mGSC displayed higher CD31 expression than pGSC as early as day 3, suggesting an accelerated endothelial-like transdifferentiation in the mesenchymal subtype. These findings indicate that the presence of endothelial cells enhances vascular mimicry in GSC organoids and that mesenchymal GSCs may possess a greater intrinsic plasticity toward endothelial lineages, likely driven by paracrine signaling and ECM-mediated interactions^65^.

Our findings demonstrate that GSC Matrigel spheroids exhibit enhanced multilineage differentiation potential when co-cultured with endothelial cells, particularly over extended culture periods. While initial exposure to Matrigel alone results in reduced differentiation, prolonged co-culture with ECs significantly enhances differentiation towards astrocytes, oligodendrocytes, pericytes, and endothelial-like cells. pGSC primarily favors astrocytic differentiation, whereas mGSC shows a broader capacity for differentiation toward oligodendrocytic, pericytic, and endothelial lineages. Importantly, both pGSC and mGSC simultaneously have increased stemness, proliferation, and hypoxia with increased culture complexity and duration, consistent with asymmetric cell division and plasticity observed in glioma stem cells. Compared to spheroids, which progressively lose their differentiation capacity, Matrigel spheroids especially under EC co-culture demonstrate a dynamic capacity to sustain stem-like features while supporting lineage diversification. These results suggest that GSC-derived Matrigel spheroids can spontaneously and progressively transition into heterocellular organoid-like structures under appropriate microenvironmental cues, offering a robust platform to study glioblastoma heterogeneity and vascular interactions in vitro.

### Development of a GBM-Relevant Engineered Extracellular Matrix (eECM)

To mimic the stiffened tumor microenvironment characteristic of recurrent glioblastoma (GBM), we designed an engineered extracellular matrix (eECM) incorporating both mechanical and biochemical features relevant to glioma stem cell (GSC) biology. Healthy brain tissue typically exhibits low stiffness (∼1 kPa), whereas GBM progression is associated with extracellular matrix (ECM) remodeling that increases local stiffness well beyond 10 kPa. In some xenograft models, stiffness values have been reported as high as 100-200 kPa, particularly in recurrent tumors. Therefore, we aimed to develop an eECM formulation with a compressive modulus in the 200-400 kPa range to model this post-treatment niche. The matrix was composed of gelatin methacrylate (GelMA) as a collagen substitute, hyaluronic acid methacrylate (HAMA) to recapitulate brain ECM components^66^, and fibronectin to support endothelial cell (EC) interactions (**Figure 4A; Materials and Methods**). All formulations contained 15 mM LAP and 1 mM tartrazine for consistent photopolymerization.

**Figure 4.**
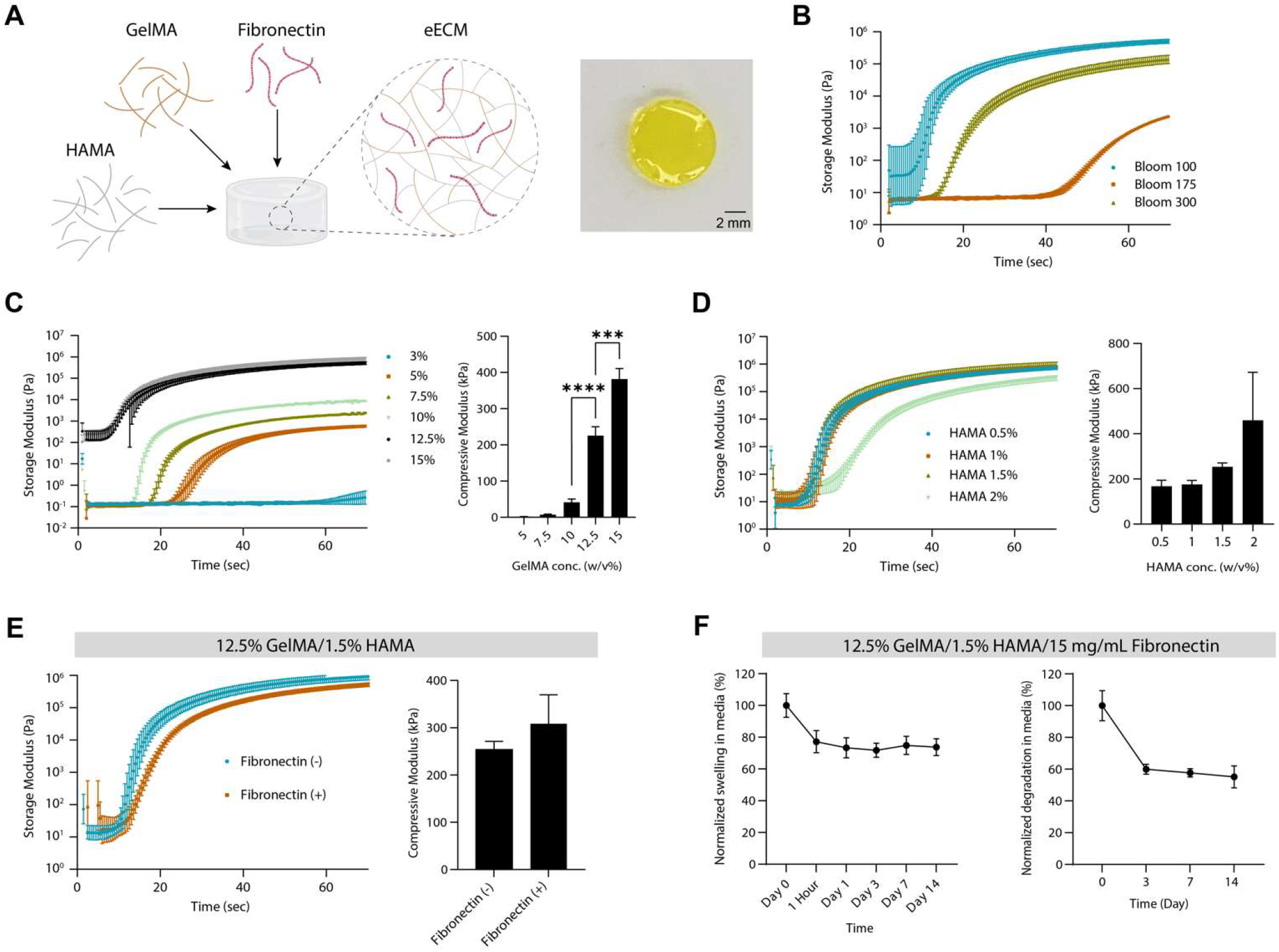
Synthesis and characterization of engineered extracellular matrix (eECM). (A) Schematic showing formulation of eECM incorporating GelMA, HAMA, and fibronectin. Representative image of crosslinked eECM disk (scale bar = 2 mm). (B) Rheological time-sweep experiment of 12.5% GelMA (bloom number 100, 175, 300) showing crosslinking kinetics when exposed to UV (11.2 mW/cm^2^, 405 nm). An increase in storage modulus (G′) with time showing initiation of crosslinking and reaching a plateau indicate fully crosslinked network. (C) Effect of increasing GelMA concentration (3–15% w/v) at fixed bloom 100 significantly increases compressive modulus (n = 3, ***p ≤ 0.0005). (D) The addition of HAMA (0.5–2% w/v) to 12.5% GelMA slightly enhances storage and compressive modulus. (E) Addition of fibronectin (15 µg/mL) to 12.5% GelMA/1.5% HAMA formulation has minimal impact on mechanical stiffness. (F) Swelling (left) and degradation (right) kinetics of final eECM formulation (12.5% GelMA/1.5% HAMA/15 µg/mL fibronectin) over 14 days in CCM+B27 at 37 °C (n = 5). Data are presented as mean ± SD. Statistical analysis was performed using two-way ANOVA with Tukey’s post hoc test (*p ≤ 0.05, **p ≤ 0.005, ***p ≤ 0.0005, ****p ≤ 0.00005).

Initial rheological analyses of 12.5% GelMA with varying bloom strengths (100, 175, and 300) revealed that GelMA Bloom 100 exhibited the highest stiffness (∼100 kPa storage modulus within 30 seconds), making it the best candidate for further optimization (**Figure 4B**). Next, we assessed GelMA Bloom 100 across a concentration range (3–15%). While lower concentrations had poor crosslinking behavior, both 12.5% and 15% achieved storage and compression moduli in the target range (**Figure 4C**). However, 15% GelMA was deemed overly rigid (∼382 kPa), whereas 12.5% GelMA (∼225 kPa) offered a more appropriate stiffness for modeling GBM recurrence and was selected as the primary hydrogel component.

To further enhance physiological relevance, we tested HAMA 10K at concentrations from 0.5% to 2% in combination with 12.5% GelMA (**Figure 4D**). Among these, only 1.5% HAMA produced compressive stiffness (∼254 kPa) within the desired 200-400 kPa range without compromising gelation dynamics. Additionally, fibronectin (15 µg/mL) was evaluated for its effects on matrix stiffness and biological function. Although its inclusion caused a slight delay in gelation and a modest increase in compressive modulus (∼309 kPa), both values remained within acceptable limits. Given fibronectin’s established role in promoting EC adhesion and angiogenesis, it was incorporated into the final eECM formulation.

Swelling and degradation studies revealed that the eECM undergoes the greatest structural changes within the first 24-72 hours of culture, with the matrix stabilizing thereafter (**Figure 4D-E**). After an initial 30% reduction in wet weight and ∼40% degradation by day 3, the material remained structurally intact through day 14. These findings suggest the eECM is suitable for short-to-intermediate term cell culture applications. In summary, the optimized eECM formulation - 12.5% GelMA Bloom 100, 1.5% HAMA 10K, and 15 µg/mL fibronectin - offers the desired mechanical properties and physiological stability for supporting GSC culture in a rigid, GBM-like environment.

### Encapsulation of GSC Matrigel Spheroids in eECM Enhances Proliferation and Differentiation in the Presence of Endothelial Cells

To evaluate the functional behavior of GSCs in a more physiologically relevant microenvironment, we encapsulated Matrigel spheroids of pGSC and mGSC in engineered extracellular matrix (eECM) alone or co-cultured them with HUVECs (**Figure 5A; Materials and Methods**). Brightfield imaging of spheroids in all five 3D culture conditions - spheroid-only, Matrigel spheroids, Matrigel + EC co-culture, eECM encapsulated, and eECM encapsulated + EC co-culture demonstrated distinct morphological evolution over time (**Figure 5B**). At day 3, spheroids across all conditions maintained compact, circular shapes. By day 14, significant differences emerged. While spheroids and Matrigel spheroids showed minor changes, co-culture with ECs led to more defined and compact spheroid boundaries. Most notably, eECM encapsulated spheroids showed denser peripheries and reduced scattering of cells into the surrounding matrix, suggesting improved architectural integrity and potentially reduced invasiveness. These changes were especially prominent in the co-culture condition, where the spheroid edges appeared darker and more structured.

**Figure 5.**
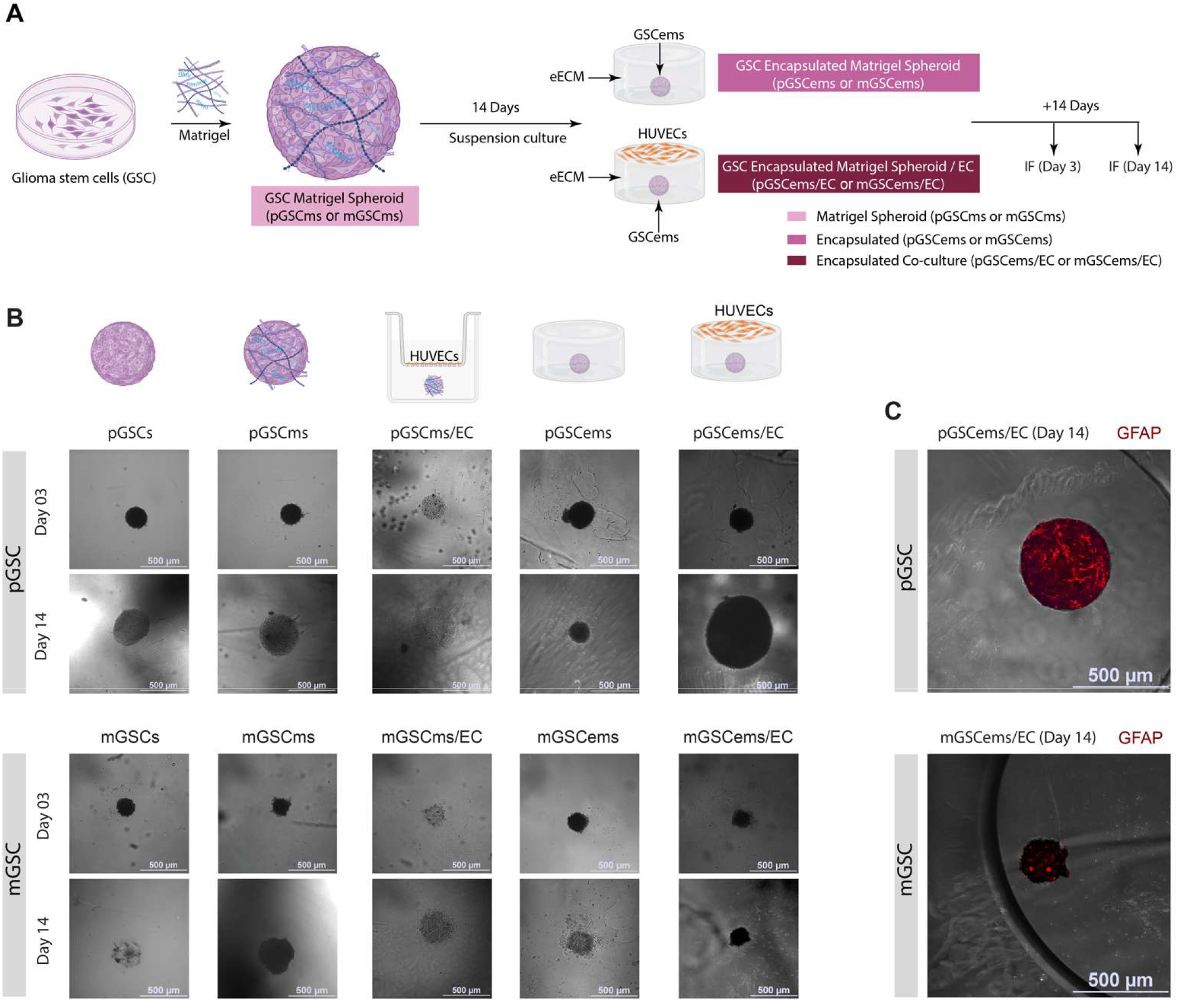
Establishing the engineered extracellular matrix (eECM) encapsulation and endothelial co-culture model. (A) Schematic representation of the experimental workflow. Glioblastoma stem cells (GSCs) were cultured as Matrigel spheroids (pGSCms/mGSCms) for 14 days, then encapsulated in engineered extracellular matrix (eECM) and maintained alone (pGSCems/mGSCems) or co-cultured with HUVECs (pGSCems/EC /mGSCems/EC) for an additional 3 or 14 days. (B) Phase contrast images of GSC grown in all 3D culture conditions (GSCs, GSCms, GSCms/EC, GSCems, and GSCems/EC) used at days 3 and 14. (C) Representative images of encapsulated pGSC and mGSC Matrigel spheroids that were co-cultured with HUVECs for 14 days before being stained for GFAP (red). Images were enhanced by overlaying the GFAP fluorescent Matrigel spheroid over the same sample’s phase contrast image showing the encapsulation.

To assess astrocytic differentiation within this platform, immunofluorescence staining for GFAP was performed on day 14 spheroids co-cultured with ECs and overlaid with corresponding phase contrast images (**Figure 5C**). Both pGSCems/EC and mGSC/EC spheroids demonstrated positive GFAP staining, with a higher signal intensity in pGSC, consistent with its proneural identity. The localization of GFAP staining primarily to the periphery of the spheroid suggests region-specific differentiation and possible radial glia-like organization. Additionally, GFAP has been shown to play a structural role in tunneling nanotubes between glioblastoma cells, thereby assisting with the intracellular transfer of mitochondria in vitro and in vivo^57^. Although some studies suggest that mesenchymal GSCs are known for their increased GFAP expression, pGSC expressed more *GFAP* than mGSC after *in vitro* culturing outside the patient (**Table S1**)^39, 50^. This pattern was less pronounced in mGSC, but detectable, indicating that eECM encapsulation combined with endothelial interaction promotes astrocytic lineage commitment across both GSC subtypes.

To further assess functional changes, we performed IF staining for markers of stemness (CD133), proliferation (Ki67), and hypoxia (CAIX) at Day 3 and Day 14 post-encapsulation (**Figure 6**, top row). CD133 levels remained relatively stable over time in both pGSC and mGSC encapsulated spheroids. Ki67 expression significantly increased on Day 14 in the presence of HUVECs, particularly in mGSC co-cultures, indicating that endothelial-derived signals promote GSC proliferation. Neural stem cells express both Ki67 and CD133 in vivo at early time points^67^. Notably, the TCGA database suggests that proneural GBM samples express more Ki67 than mesenchymal samples, however the CGGA database suggests that these subtypes have similar expression levels in vivo (**Figure S2B**)^53, 54^. CAIX staining showed minimal hypoxic response across all conditions, suggesting that the encapsulated spheroids were adequately oxygenated, or that hypoxia develops later in culture. Although the GBM patients from TCGA suggest similar CAIX expression levels for both subtypes, the GBM patients from CGGA have more CAIX expression levels in mesenchymal subtype (**Figure S2C**)^53, 54^. Altogether, these results indicate that the encapsulation model developed herein recapitulates the GSC health expression patterns observed in patients.

**Figure 6.**
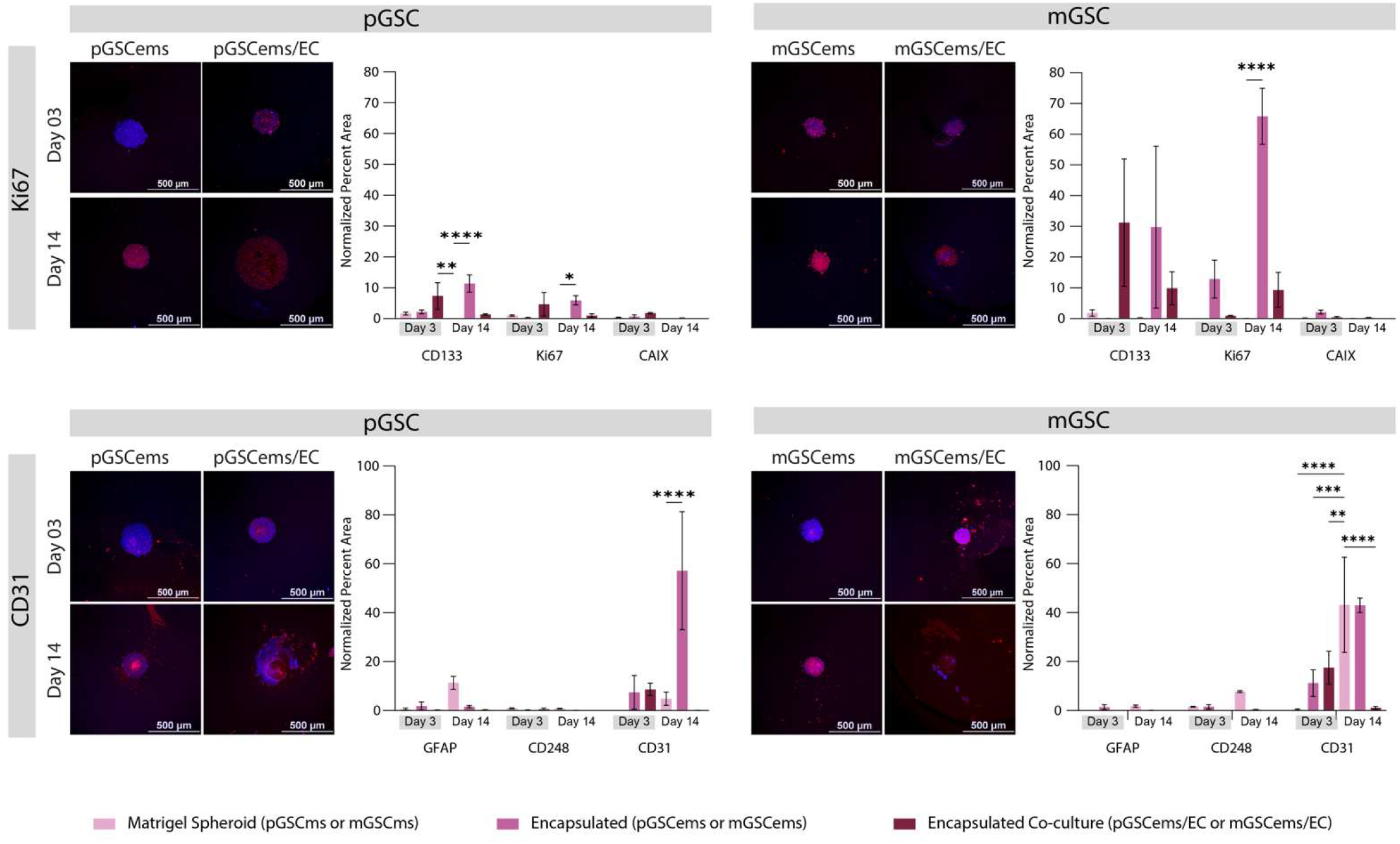
Engineered extracellular matrix (eECM) encapsulation and endothelial co-culture promote GSC differentiation, stemness loss. Top row displays immunofluorescence (IF) staining and quantification of stemness (CD133), proliferation (Ki67), and hypoxia (CAIX) markers in pGSC and mGSC spheroids under encapsulated and co-culture conditions. Bottom row shows IF staining and quantification of astrocytic (GFAP), oligodendrocytic (MOG), pericytic (CD248), and endothelial (CD31) differentiation markers, highlighting enhanced lineage specification with EC co-culture. Data are presented as mean ± SD (N = 3). Statistical analysis was performed using two-way ANOVA with Tukey’s post hoc test (*p ≤ 0.05, **p ≤ 0.005, ***p ≤ 0.0005, ****p ≤ 0.00005).

We next analyzed the expression of differentiation markers for astrocytic (GFAP), pericytic (CD248), and endothelial (CD31) lineages (**Figure 6**, bottom row). Quantification revealed that co-culture with HUVECs significantly enhanced differentiation in both GSC subtypes, with pGSC showing higher GFAP expression and mGSC exhibiting stronger expression of CD248, and CD31. These results are consistent with subtype-specific lineage bias, where pGSC preferentially differentiates toward astrocytes and mGSC toward pericytes, and endothelial-like cells^65, 68, 69^. Importantly, endothelial co-culture not only promoted higher expression of these genes but also accelerated the acquisition of these phenotypes over time. GFAP is similarly expressed in both subtypes found in patient samples (**Figure S2D**)^53, 54^. CD248 expressed by more mesenchymal patient samples than proneural patient samples (**Figure S2F**)^53, 54^. Although no CD31 (PECAM1) data is available within the CGGA dataset, the TCGA dataset indicates that mesenchymal patient samples express more CD31 than proneural samples (**Figure S2G**)^53, 54^. Taen together, these findings reflect previously known expression patterns, highlight the potential for GSCs to differentiate towards vascular-supportive cell-types, and emphasize the ability of our novel platform to quickly adapt towards a perivascular niche state in a recurrent-like environment.

Together, these findings highlight the utility of the eECM encapsulation platform combined with endothelial co-culture to model GSC behavior in a complex, biomimetic environment. This system supports both proliferation and lineage-specific differentiation, offering a powerful tool to investigate GBM progression and therapeutic responses in vitro.

## DISCUSSION

Numerous studies have demonstrated that patient-derived cancer cells exhibit markedly different gene expression profiles when cultured in traditional 2D monolayers compared to 3D spheroidal systems, underscoring the need for more physiologically relevant in vitro models of GBM^70-72^. While 3D culture platforms using Matrigel have advanced the field, we still have several limitations^73-75^. Our findings corroborate prior reports that proliferation, as measured by Ki67 expression, decreases in Matrigel-only environments but is restored when glioma stem cells (GSCs) are co-cultured with endothelial cells (ECs)^26, 73, 75^.

The Matrigel spheroid platform developed in this study supports concurrent increases in stemness, proliferation, and differentiation markers with time and increasing microenvironmental complexity. This dual increase suggests that the spheroids undergo asymmetric cell division, a hallmark of cancer stem cell behavior, including that of GSCs^76, 77^. The ability of this system to maintain a stem-like population while also generating lineage-committed cells offers a powerful platform to mimic tumor heterogeneity and study cellular dynamics in a controlled 3D context. Additionally, our Matrigel spheroids are robust enough to withstand transferring to a different location^22^.

In terms of mechanical microenvironment, our results align with literature reporting that healthy brain tissue, particularly the cerebellum, where GBM often arises, has a baseline stiffness around 1 kPa, rarely exceeding 7 kPa^78, 79, 80^. GBM tumors, however, remodel their local matrix to significantly increase stiffness, particularly at the invasive edge, often surpassing 10 kPa^80, 81, 82^. Xenograft models further demonstrate ECM remodeling by GBM subtypes, with U87MG reaching ∼27 kPa and U251 models exceeding 100 kPa^83, 84^. GSCs also display increased migratory behavior on rigid substrates, with optimal motility reported at stiffness levels >200 kPa^84, 85^. Moreover, GBM recurrence occurs in an even stiffer post-treatment environment, where residual GSCs differentiate into various tumor-supportive phenotypes^86, 87^.

To address this post-treatment context, we developed an engineered ECM (eECM)-based model that recapitulates key features of GBM recurrence. Our approach begins with GSC Matrigel spheroids, which progressively self-organize into organoid-like structures via asymmetric division. This transition supports the maintenance of a GSC subpopulation alongside newly differentiated tumor-supportive lineages. Notably, in the proneural pGSC line, we observe co-expression of Ki67 and CD133, known to characterize early neural stem cells in vivo^88^. Likewise, the mGSC mesenchymal line exhibits robust CD248 expression in eECM-encapsulated co-culture, closely reflecting in vivo mesenchymal tissue features^69^.

The developed eECM not only mimics the rigid, fibrotic microenvironment of recurrent GBM but also supports long-term spheroid stability and biologically relevant differentiation patterns. While the stiffness used in this study was tailored to model recurrence-associated rigidity, our system is modular: hydrogel component concentrations and crosslinking times can be readily adjusted to simulate other mechanical contexts. Future applications may leverage this eECM for bioprinting to explore GBM mechanobiology, heterogeneity, and therapeutic resistance in highly customizable formats.

## CONCLUSION

In summary, we developed a modular and physiologically relevant glioblastoma model that integrates patient-derived GSCs, endothelial cells, and a tunable engineered extracellular matrix to recapitulate critical features of the GBM perivascular niche and recurrence-associated rigidity. Our findings demonstrate that Matrigel spheroids facilitate the transition of GSCs into organoid-like structures capable of maintaining stemness while acquiring lineage-specific differentiation potential. Endothelial co-culture significantly enhances stemness, proliferation, hypoxia, and multilineage commitment across proneural and mesenchymal GSC subtypes. Furthermore, encapsulation within a high-stiffness eECM mimicking the recurrent tumor microenvironment promotes subtype-specific phenotypes and enables long-term structural integrity. This vascularized and mechanically tunable platform offers a powerful tool for investigating GBM heterogeneity, tumor-vascular interactions, and therapeutic responses, advancing preclinical modeling for both mechanistic studies and precision oncology applications.

## MATERIALS AND METHODS

### Cell Culture

All cell cultures were maintained at 37 °C in a humidified incubator with 5% CO_2_. Mycoplasma contamination was routinely monitored using a PCR-based Mycoplasma detection kit (Thermo Fisher Scientific, Cat. No. J66117.AMJ), and all cultures tested negative. Adherent monolayer cells were passaged using Accutase (STEMCELL Technologies, Cat. No. 07922) following the manufacturer’s protocol. Live cells were centrifuged at 300 × g for 5 minutes unless otherwise specified.

### Glioma Stem Cell Culture

Patient-derived glioma stem cell (GSC) lines 3028 (pGSC) and 3128 (mGSC) were generously provided by Baylor College of Medicine (Houston, TX). Both cell lines were originally derived from xenografted human GBM tumors, dissociated, and selected for CD133+ expression. pGSC and mGSC were subsequently classified as belonging to the proneural and mesenchymal GBM subtypes, respectively, and were stably transduced with GFP via lentiviral vectors prior to transfer. GSCs were cultured in Neurobasal-A medium (Thermo Fisher Scientific, Cat. No. 12349015) supplemented with the following final concentrations: 1x GlutaMAX (from 100x stock, Cat. No. 35050061), 1x sodium pyruvate (from 100 mM stock, Cat. No. 11360070), 1x antibiotic-antimycotic (from 100x stock, Cat. No. 15240096), 20 ng/mL human EGF recombinant protein (Cat. No. PHG0311L), 20 ng/mL human FGF-basic recombinant protein (Cat. No. PHG0261), and 2x B-27™ Supplement minus vitamin A (from 50x stock, Cat. No. 12587010). A 2% Matrigel coating solution was prepared by diluting 1 mL of Corning® Matrigel® hESC-Qualified Matrix (Corning, Cat. No. 356278) into 49 mL of ice-cold 1x PBS. T-25 or T-75 flasks were coated on one surface with the Matrigel solution and incubated at 37 °C and 5% CO_2_ for 1 hour prior to cell seeding. GSCs were seeded at 1 × 10^5^ cells/mL and cultured adherently on the coated surface until reaching approximately 80% confluence. Cells were passaged up to passage 11 and maintained in monolayer culture with media changes every 2-3 days. To confirm retention of stem-like properties, neurosphere formation assays were performed every few passages using non-adherent culture conditions.

### Endothelial Cell Culture

Human umbilical vein endothelial cells (HUVECs) were obtained from multiple donors (PromoCell GmbH, Cat. No. C-12203). Cells were cultured in Endothelial Cell Medium (ScienCell, Cat. No. 1001) supplemented with fetal bovine serum (FBS; ScienCell, Cat. No. 0025), Endothelial Cell Growth Supplement (ECGS; ScienCell, Cat. No. 1052), and penicillin/streptomycin (ScienCell, Cat. No. 0503). For flask preparation, a 1 mg/mL fibronectin stock solution was prepared by dissolving 5 mg of fibronectin (Thermo Fisher Scientific, Cat. No. 33016015) in 5 mL of autoclaved distilled water. A working 2% fibronectin coating solution was prepared by diluting 200 µL of the stock into 10 mL of 1x PBS. Culture flasks were coated on one surface with the fibronectin solution and incubated at 37 °C with 5% CO_2_ for 2 hours. HUVECs were seeded at a density of 1 × 10^5^ cells/mL and cultured as adherent monolayers on the coated surface until reaching ∼80% confluence. Cells were maintained up to passage 15, with media changes every 2–3 days.

### Co-culture

Co-culture medium was prepared using Endothelial Cell Basal Medium (PromoCell, Cat. No. C-22210) supplemented with the following components: 1× GlutaMAX (from 100x stock; Thermo Fisher Scientific, Cat. No. 35050061), 1x antibiotic-antimycotic (from 100x stock; Thermo Fisher Scientific, Cat. No. 15240096), 2x B-27™ Supplement minus vitamin A (from 50x stock; Thermo Fisher Scientific, Cat. No. 12587010), 20 ng/mL human EGF recombinant protein (Cat. No. PHG0311L), 20 ng/mL human FGF-basic recombinant protein (Cat. No. PHG0261), 0.2 µg/mL hydrocortisone (from 50 µg/mL stock; PromoCell, Cat. No. C-39211), 0.4% endothelial cell growth supplement (PromoCell, Cat. No. C-39210), and 0.5 ng/mL VEGF 165 (from 10 µg/mL stock; PromoCell, Cat. No. C-39211). For co-culture setup, surfaces intended for HUVEC seeding were first coated with 2% fibronectin solution and incubated at 37 °C with 5% CO_2_ for 20 minutes. HUVECs were then seeded onto the coated surfaces at a density of 2 × 10^6^ cells/mL.

### Spheroid and Matrigel Spheroid Formation

GSC spheroids and Matrigel spheroids were generated in BRANDplates® inertGrade™ U-bottom 96-well microplates (BrandTech Scientific, Cat. No. 781900). Wells were seeded with 100 µL of GSC suspension in GSC medium at a concentration of 1 × 10^4^ cells/mL for immunofluorescence (IF) samples or 1 × 10^5^ cells/mL for qRT-PCR samples. Plates were centrifuged at 300 x g for 5 minutes and incubated at 37 °C with 5% CO_2_ on a plate shaker set to ∼60 rpm. For Matrigel spheroids, the protocol was identical to spheroid formation through the initial centrifugation step. Following centrifugation, 4 µL of 2% Matrigel solution was added to each well, and the plates were centrifuged again for 5 minutes at 300 x g. Plates were then returned to the incubator on the plate shaker at 37 °C and 5% CO_2_.

### Spheroid and Matrigel Spheroid Maintenance

All spheroids were continuously cultured on a plate shaker at 37 °C and 5% CO_2_. On day 3, an additional 100 µL of GSC media was added to each well. Culture volume was maintained between 100–200 µL to account for evaporation (notably in outer wells). Media was refreshed once per week, with careful handling to avoid accidental loss of spheroids. Under these conditions, both spheroids and Matrigel spheroids were successfully maintained for up to two months.

### Transwell® Co-culture with Matrigel Spheroids

The day prior to co-culture, Transwell® 96-well insert plates were pre-coated with 20 µg/mL fibronectin at room temperature for at least 15 minutes. The receiver plate wells were filled with 300 µL of HUVEC medium. HUVECs were seeded onto the insert membranes at a density of 2 × 10^6^ cells/mL and incubated overnight at 37 °C. On the day of co-culture, two-week-old Matrigel spheroids had their medium replaced with 200 µL of co-culture medium per well. HUVEC medium was carefully aspirated from the insert wells to avoid disrupting the adherent cells. The insert plate was then placed into the spheroid plate, and additional co-culture medium was added to the insert wells. Transwell® co-culture plates were returned to the incubator on a plate shaker at ∼60 rpm and maintained for either 3 or 14 days, depending on the experimental condition.

### In Situ Encapsulation of Matrigel Spheroids

Matrigel spheroids were cultured for 14 days as described previously. To begin encapsulation, culture medium was carefully aspirated, leaving approximately 5 µL of residual medium in each well to prevent spheroid loss. Next, 50 µL of filter-sterilized, pre-warmed (37 °C) hydrogel-based extracellular matrix (hbECM) was gently added to each well. Plates were centrifuged at 300 × g for 5 minutes to center the spheroids within the gel. Encapsulation was achieved by crosslinking each well individually using an OmniCure system equipped with a 405 nm filter. Wells were exposed to light at ∼11.2 W/m^2^ for 1 minute at a distance of ∼6 cm. Following crosslinking, samples were gently rinsed by incubating in 1× PBS for 5 minutes. For single-culture conditions, PBS was aspirated and replaced with 100 µL of cNBA + B-27 medium. For co-culture conditions, the PBS was first removed, and the hydrogel surface was coated with 20 µg/mL fibronectin for at least 15 minutes at room temperature. Subsequently, 100 µL of HUVEC suspension (2 × 10^6^ cells/mL in CCM + B-27 medium) was added to the surface. All cultures were maintained at 37 °C and 5% CO_2_ with daily media changes for the first 3 days, followed by changes every 2-3 days for the duration of the experiment.

### Synthesis of engineered extracellular matrix (eECM)

Gelatin methacryloyl (GelMA) was synthesized as previously described^89^. Briefly, 10 g of gelatin from porcine skin (type A, gel strength 80-120 g bloom; Sigma-Aldrich, Cat. No. G6144-100G) was dissolved in 100 mL of 1x PBS at 50 °C under constant stirring. The pH of the solution was adjusted to approximately 8.5 using 1 N NaOH added dropwise. To achieve a high degree of methacrylation, 8 mL of methacrylic anhydride (Alfa Aesar, Cat. No. A13030) was added dropwise and allowed to react for 3 hours at 50 °C. The reaction mixture was centrifuged at 1,438 x g for 5 minutes at 28 °C. The supernatant was collected and diluted in 400 mL of pre-warmed (37 °C) 1x PBS to quench the reaction. The resulting GelMA solution was dialyzed against deionized water (DiH_2_O) at 50 °C for 7 days using 12-14 kDa MWCO dialysis tubing, with water changes 2-3 times per day. The dialyzed product was frozen at −80 °C overnight and lyophilized for 5 days. Variants GelMA B175 and B300 were prepared using the same procedure.

Hyaluronic acid methacrylate (HAMA) was synthesized as previously reported^90^. In brief, 5 g of hyaluronic acid (MW 10 kDa; HAWORKS, Cat. No. HA-10KXia) was dissolved in 100 mL of DiH_2_O in an ice bath with constant stirring overnight. While maintaining the temperature in the ice bath, 30 mL of methacrylic anhydride was added dropwise. The pH was adjusted to 8.5-9.5 using 1 N NaOH and monitored every 30 minutes for 6 hours, with additional NaOH added as needed to maintain the pH. The reaction was allowed to proceed overnight with vigorous stirring in the ice bath. The reaction mixture was then centrifuged at 1,438 x g for 5 minutes at 4 °C. The supernatant was dialyzed in DiH_2_O at room temperature for 7 days using 12-14 kDa MWCO tubing, with water changes 2-3 times daily. The dialyzed solution was frozen at −80 °C overnight and lyophilized for 5 days.

To prepare the engineered extracellular matrix (eECM), concentrated stock solutions were prepared as follows: 25% w/v GelMA B100 HS, 10% w/v HAMA 10K, 100 mM lithium phenyl-2,4,6-trimethylbenzoylphosphinate (LAP; CAS 85073-19-4; AmBeed, Cat. No. A636763), 100 mM tartrazine (CAS 1934-21-0; TargetMol, Cat. No. TN2258), and 1 mg/mL fibronectin (Life Technologies, Cat. No. 33016015). All components were pre-warmed to 37 °C and then mixed to achieve final concentrations of 12.5% w/v GelMA B100 HS, 1.5% w/v HAMA 10K, 15 mM LAP, 1 mM tartrazine, and 15 µg/mL fibronectin. The mixture was vortexed thoroughly, briefly heated (∼1 minute), and sterile-filtered using a Millex-GV Durapore PVDF membrane syringe filter (0.22 µm pore size, 33 mm diameter; Cat. No. SLGVR33RB).

### Characterization of eECM

To assess the degradation and swelling behavior of the engineered extracellular matrix (eECM), samples were prepared and crosslinked under standardized conditions (λ = 405 nm, distance = 6 cm, intensity = 11.2 mW/cm^2^, exposure time = 1 minute) in the absence of cells. For degradation analysis, crosslinked eECM hydrogels were incubated in complete co-culture medium (CCM + B27) at 37 °C for 3, 7, or 14 days. At each time point, samples were freeze-dried, and the dry weights were recorded. The initial dry weight of eECM (day 0) was determined by immediately freeze-dried the scaffold following crosslinking without incubation. Degradation was quantified by calculating the normalized dry weight as follows: Normalized Weight (%) = (Dry weight at Day x) / (Initial dry weight at Day 0) x 100%. For swelling analysis, two experimental approaches were used. First, to evaluate swelling from a lyophilized state, crosslinked and freeze-dried eECM samples were weighed (Initial dry weight at 0 minute), then incubated in complete media (CCM + B27) at 37 °C. Samples were removed at 5-minute intervals for 1 hour, blotted with filter paper, and weighed to determine wet mass. Swelling was calculated as: Normalized Weight (%) = (Minute x / Initial dry weight at 0 minute) × 100%. Second, to assess swelling under in situ conditions (i.e., post-crosslinking but without lyophilization), eECM samples were crosslinked and incubated directly in complete medica (CCM + B27) at 37 °C for 1 hour, 1 day, 3 days, 7 days, or 14 days. Initial wet weight was weighed immediately after crosslinking. At each time point, samples were blotted and weighed, and swelling was calculated as: Normalized Weight (%) = (Wet weight at time x) / (Initial wet weight) × 100%.

The viscoelastic properties of the hydrogel ink were evaluated using a Discovery HR-20 hybrid rheometer (TA Instruments, USA) equipped with a Peltier temperature control system and a 20 mm stainless steel parallel plate geometry set at a 1 mm gap. Time sweep measurements were conducted at 25°C for 70 seconds under oscillatory conditions (10 rad/s, 1% strain) within the linear viscoelastic region. Crosslinking was initiated by exposure to 405 nm UV light (11.2 mW/cm^2^), and the evolution of the storage modulus (G′) and loss modulus (G′′) was monitored in real time. Formulations containing varying concentrations of GelMA, HAMA, and fibronectin were tested, while the photoinitiator (LAP, 15 mM) and photoabsorber (tartrazine, 1 mM) were kept constant.

For mechanical testing, swollen hydrogel disks (8 mm diameter, 1.5–2 mm thickness) were subjected to uniaxial compression using an ADMET Expert 7600 series testing system at room temperature (23 ± 2 °C). Samples were compressed at a constant rate of 1 mm/min, and the Young’s modulus (E) was calculated from the slope of the linear region of the stress-strain curve up to 30% strain.

### Encapsulation of GSC Spheroids within eECM

Matrigel spheroids were first cultured for 14 days as previously described. To encapsulate them in eECM, media was gently aspirated from each well, leaving approximately ∼5 µL to prevent loss of spheroids. A minimum of 20 µL of filter-sterilized, pre-warmed (37 °C) eECM was then added to each well. Spheroids suspended in eECM were carefully transferred to the desired location using a 200 µL pipette or larger to minimize shear stress and preserve spheroid morphology.

### RNA Purification

To purify RNA from spheroid or matrigel spheroid (including co-culture), samples from at least three 96well plates of the same strain, passage, condition, and time were collected and combined into one 15 mL centrifuge tube. This involved using a 1000 µL pipette to transfer all media and samples from each well till the tube was filled. Once filled, the tube was centrifuged for 5 minutes at 300 x g and the media aspirated. This process repeats until all 288 spheroids/matrigel spheroids have been collected. When all samples are collected, pelleted, and media aspirated, samples were resuspended in 1 mL Accutase® and incubated overnight (∼12-16 hrs) at 4°C. The following morning, samples were vortexed on high for 30 seconds or until all spheroidal samples were disrupted into a single cell state. The tube was then centrifuged for 5 minutes at 300g and the Accutase® aspirated. Cells were briefly resuspended in sterile 1x PBS to count the cells. After counting cells, approximately 1-2×10^6^ cells were isolated, pelleted, and resuspended in 600 µL TRIZOL® (Zymo Research, R2050-1-200) method. Concentration of the isolated RNA was determined with NanoDrop. A total of 1μg of RNA was then converted into cDNA using qScript™ cDNA SuperMix (QuantaBio Cat # 101414-106).

### Gene expression Analysis

Quantitative real-time polymerase chain reaction (qRT-PCR) was performed using comparative Ct (ΔΔCt) experiments on the QuantStudio™ 3 System (Applied Biosystems). Final concentration of 12.5 ng RNA template was used for qPCR using 1x PowerUp™ SYBR™ Green Master Mix (ThermoFisher Scientific, Cat # A25742). GAPDH served as the internal control for mRNA expression normalization.

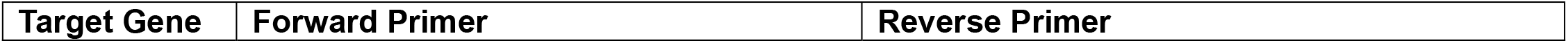
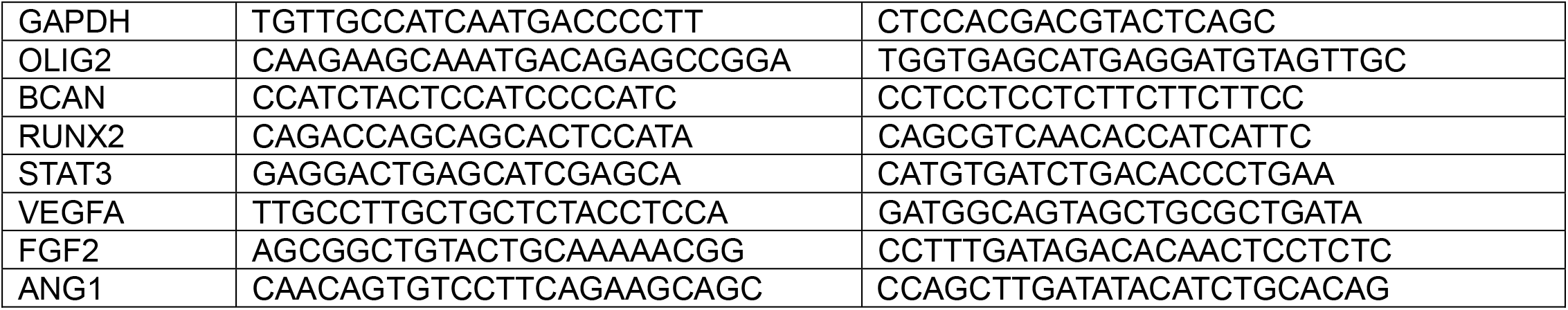

### Immunofluorescent Staining

To prepare samples for immunofluorescent (IF) staining, media was first removed and then blocked and permeabilized with 5% FBS and 0.1% Triton X-100 in PBS while protected from light. Next, samples were washed three times in 1x PBS. After washing, samples were incubated in primary antibody (1:100 dilution in antibody dilution buffer: 1% BSA and 0.3% Triton X-100 in 1x PBS)/well at 4°C overnight (12-24 hr). Samples were again washed three times in 1x PBS followed by incubation in secondary antibody (1:100 dilution in antibody dilution buffer). Samples were again washed for three times in 1x PBS followed by incubation in DAPI (1:10,000 dilution of 10.9 mM stock DAPI in 1x PBS) (Fisher Scientific, Cat # D3571) followed by a final round of three 1x PBS washes. Non-encapsulated samples were then stored in Fluoro-Gel (with Tris Buffer) (Electron Microscopy, Cat # 17985-10). Encapsulated samples were stored in 1x PBS. All antibodies were purchased from ProteinTech (CD133 - Cat # 18470-1-AP, Ki67 – Cat # 27309-1-AP, CAIX - Cat # 11071-1-AP, GFAP - Cat # 23935-1-AP, MOG - Cat # 12690-1-AP, CD248 (TEM1) - Cat # 60170-1-Ig, CD31 - Cat # 11265-1-AP, Coralite 594-conjugated mouse anti-heavy chain rabbit IgG - Cat # SA00014-5, CoraLite 594-conjugated rabbit anti-heavy chain mouse IgG – Cat # CL594-10283).

| IF Staining Step | Non-encapsulated | Encapsulated |
| --- | --- | --- |
| Block and permeabilize | 30 minutes, room temperature | 12 hours, 4°C |
| 1x PBS washes | 5 minutes, room temperature | 30 minutes, room temperature |
| Primary antibody | 12 hours, 4°C | 24 hours, 4°C |
| Secondary antibody | 1 hour, room temperature | 6 hours, room temperature |
| DAPI | 15 minutes, room temperature | 2 hours, room temperature |

Samples were imaged using Nikon Eclipse Ti confocal microscope with 25 µm z-steps to capture full spherical range with a 10x objective. Images were processed using NIS Elements software version 4.0 and ImageJ version 1.54p. Thumbnails are the maximum intensity projections. Intensities were quantified by first creating a sum intensity projection of the DAPI and TRITC layers. Then a mask was created from the DAPI layer such that only the area of cellular presence would be quantified. That area mask was then applied to the TRITC layer, dilated once, and the threshold was set to positive signal levels (see table below). The final quantification is equal to the percentage of the area of positive TRITC signal in relation to the positive DAPI signal.

| Marker | Minimum Intensity | Maximum Intensity | Particle Size |
| --- | --- | --- | --- |
| CD133 | 735448 | 1.00E+30 | 0 - Infinity |
| Ki67 | 1.19E+06 | 1.00E+30 | 0 - Infinity |
| CAIX | 965419 | 1.00E+30 | 0 - Infinity |
| GFAP | 1492940 | 1.00E+30 | 0 - Infinity |
| MOG | 348802 | 1.00E+30 | 0 - Infinity |
| CD248 | 803108 | 1.00E+30 | 0 - Infinity |
| CD31 | 624492 | 1.00E+30 | 0 - Infinity |
| DAPI | whatever fills spheroid edges with minimal external noise | 1.00E+30 | 500 - Infinity |

## STATISTICAL ANALYSIS

Quantitative data are presented as mean ± standard deviation (SD). Statistical analyses were performed using GraphPad Prism (v10.5.0). Statistical significance was evaluated using either an unpaired t-test or one-way or two-way analysis of variance (ANOVA), followed by Tukey’s post hoc multiple comparison test, with *p ≤ 0.05, **p ≤ 0.005, ***p ≤ 0.0005, ****p ≤ 0.00005 considered significant.

## AUTHOR CONTRIBUTIONS

Conceptualization and Funding Acquisition: A.G., I.S.

Experimental Design: A.M.

Cell Culture: A.M., C.J.

RNA Purification: A.M., C.J., S.M.

qRT-PCR Experiments: A.M., C.J.

qRT-PCR Analysis: A.M., C.J., A.S.

Immunofluorescent Staining and Imaging: A.M.

Immunofluorescent Analysis: A.M., C.J., A.S., R.S., M.M.

Biomaterials Development and Characterization: A.M., R.S.

Writing – Original Draft: A.M., C.J., A.S., A.G., I.S.

Writing – Review & Editing: A.M., A.G., I.S.

## ACKNOWLEDGEMENTS

The authors gratefully acknowledge Dr. Charles Lin from Baylor College of Medicine for providing us with pGSC and mGSC. We also thank Dr. Thuy-Uyen Nguyen, Dr. Min Hee Kim and Biswadeep Nayak for their initial assistance with cell culture and biomaterials development. We would also like to acknowledge the use of Texas A&M University Microscopy and Imaging Center Core Facility (RRID:SCR_022128) for confocal microscopy. The results shown here are in whole or part based upon data generated by the TCGA Research Network: https://www.cancer.gov/tcga, specifically using the CGGA and TCGA-GBM databases. All figures were organized in Adobe Illustrator. The authors used AI-based language editing tools solely to improve grammar, readability, and clarity. All scientific content and conclusions were developed, written, and verified by the authors.

## Funding

National Institute of Neurological Disorders and Stroke (NINDS) (R21 NS121945)

National Institute of Dental and Craniofacial Research (NIDCR) R01 DE032031 (AKG, IS)

National Cancer Institutes (NCI) R01 CA282251 (IS)

Cancer Prevention and Research Institute of Texas (CPRIT) RP230204 (IS)

Texas A&M University Health Science Center Seedling Grant (IS, AKG)

The content is solely the responsibility of the authors and does not necessarily represent the official views of the funding agencies.

## Data access statement

The data supporting the findings of this study are available at the Zenodo repository: 10.5281/zenodo.16907365

## Ethics statement

This study did not involve human participants or live vertebrate animals. All experimental protocols involving human cell lines were approved by the Institutional Biosafety Committee (IBC) at Texas A&M University and conducted in accordance with institutional guidelines and biosafety regulations.

