## Supplemental Figures for "Understanding proneural–mesenchymal transition using patient-derived glioma stem-like cell (GSC) organoids and engineered extracellular matrix"

**Supplementary Table S1. Gene expression levels from Mack, et al, *Chromatin landscapes reveal developmentally encoded transcriptional states that define human glioblastoma*, 2019[39].** Gene expression levels in Fragments Per Kilobase of transcript per Million mapped reads (FPKM) of GSCs. Reference names from the 2019 paper in parentheses after nomenclature used in this study. Selected results are markers for proneural GSCs (OLIG2, BCAN), mesenchymal GSCs (STAT3, RUNX2), angiogenesis (VEGFA, FGF2), stemness (CD133), proliferation (Ki67), hypoxia (CAIX), astrocytic differentiation (GFAP), oligodendrocytic differentiation (MOG), pericytic differentiation (CD248), and endothelial differentiation (CD31).

| GENE Symbol | pGSC (GSC1) | mGSC (GSC17) |
|---|---|---|
| ***OLIG2*** | 35.56151990 | 0.04018502 |
| ***BCAN*** | 12.82729650 | 2.07809460 |
| ***STAT3*** | 44.98286630 | 101.00168300 |
| ***RUNX2*** | 0.01833006 | 4.81458184 |
| ***VEGFA*** | 49.95237240 | 19.14081730 |
| ***FGF2*** | 4.68251591 | 2.20753411 |
| ***CD133 (PROM1)*** | 2.31047632 | 19.56750560 |
| ***Ki67 (MKI67)*** | 7.74293101 | 17.65308010 |
| ***CAIX (CA9)*** | 0.63039461 | 0.18397750 |
| ***GFAP*** | 8.97649805 | 0.38562261 |
| ***MOG*** | 0 | 0.00892049 |
| ***CD248*** | 0 | 0.06870684 |
| ***CD31 (PECAM1)*** | 0 | 0.36166517 |

| TCGA | CGGA |
|---|---|

**A** CD133

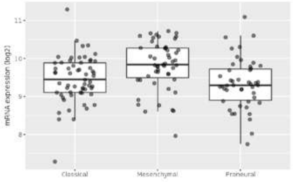


| | Sample_count | median | mad | mean | sd |
|---|---|---|---|---|---|
| Classical | 59 (37.82%) | 9.44 | 0.55 | 9.45 | 0.61 |
| Mesenchymal | 51 (32.69%) | 9.84 | 0.60 | 9.81 | 0.64 |
| Proneural | 46 (29.49%) | 9.30 | 0.63 | 9.34 | 0.66 |
| TOTAL | 156 | 9.59 | 0.60 | 9.53 | 0.66 |

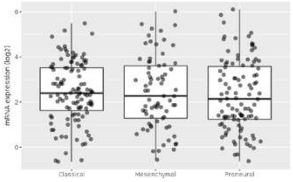


| | Sample_count | median | mad | mean | sd |
|---|---|---|---|---|---|
| Classical | 111 (38.68%) | 2.41 | 1.52 | 2.44 | 1.34 |
| Mesenchymal | 73 (25.44%) | 2.26 | 1.82 | 2.43 | 1.60 |
| Proneural | 103 (35.89%) | 2.13 | 1.68 | 2.30 | 1.57 |
| TOTAL | 287 | 2.31 | 1.65 | 2.38 | 1.49 |

**B** Ki67

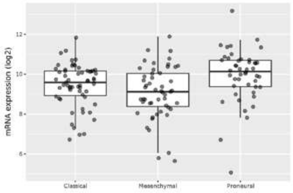


| | Sample_count | median | mad | mean | sd |
|---|---|---|---|---|---|
| Classical | 59 (37.82%) | 9.58 | 0.87 | 9.44 | 1.08 |
| Mesenchymal | 51 (32.69%) | 9.12 | 1.29 | 9.07 | 1.33 |
| Proneural | 46 (29.49%) | 10.14 | 0.94 | 9.93 | 1.35 |
| TOTAL | 156 | 9.65 | 1.15 | 9.46 | 1.28 |

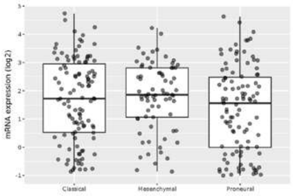


| | Sample_count | median | mad | mean | sd |
|---|---|---|---|---|---|
| Classical | 111 (38.68%) | 1.73 | 1.80 | 1.69 | 1.46 |
| Mesenchymal | 73 (25.44%) | 1.86 | 1.24 | 1.80 | 1.20 |
| Proneural | 103 (35.89%) | 1.56 | 1.81 | 1.35 | 1.52 |
| TOTAL | 287 | 1.72 | 1.62 | 1.60 | 1.43 |

**C** CAIX

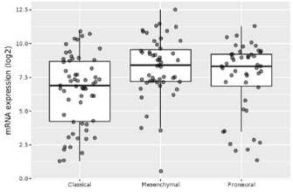


| | Sample_count | median | mad | mean | sd |
|---|---|---|---|---|---|
| Classical | 59 (37.82%) | 6.90 | 2.90 | 6.53 | 2.62 |
| Mesenchymal | 51 (32.69%) | 8.41 | 1.82 | 8.27 | 2.21 |
| Proneural | 46 (29.49%) | 8.32 | 1.64 | 7.51 | 2.62 |
| TOTAL | 156 | 7.66 | 2.26 | 7.39 | 2.58 |

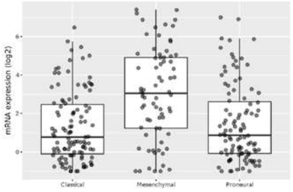


| | Sample_count | median | mad | mean | sd |
|---|---|---|---|---|---|
| Classical | 111 (38.68%) | 0.77 | 1.73 | 1.30 | 1.81 |
| Mesenchymal | 73 (25.44%) | 3.04 | 2.77 | 3.05 | 2.41 |
| Proneural | 103 (35.89%) | 0.86 | 1.73 | 1.47 | 2.01 |
| TOTAL | 287 | 1.26 | 2.12 | 1.80 | 2.17 |

**D** GFAP

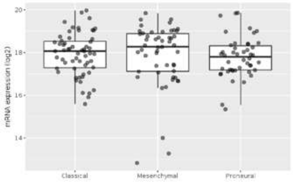


| | Sample_count | median | mad | mean | sd |
|---|---|---|---|---|---|
| Classical | 59 (37.82%) | 18.06 | 0.91 | 17.92 | 0.99 |
| Mesenchymal | 51 (32.69%) | 18.26 | 1.13 | 17.88 | 1.47 |
| Proneural | 46 (29.49%) | 17.80 | 0.90 | 17.78 | 0.95 |
| TOTAL | 156 | 18.01 | 1.09 | 17.87 | 1.15 |

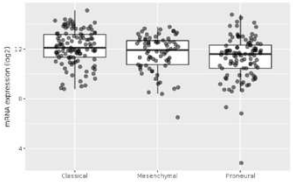


| | Sample_count | median | mad | mean | sd |
|---|---|---|---|---|---|
| Classical | 111 (38.68%) | 12.11 | 1.39 | 12.13 | 1.39 |
| Mesenchymal | 73 (25.44%) | 11.91 | 1.24 | 11.64 | 1.46 |
| Proneural | 103 (35.89%) | 11.59 | 1.35 | 11.29 | 1.74 |
| TOTAL | 287 | 11.85 | 1.47 | 11.70 | 1.58 |

**E** MOG

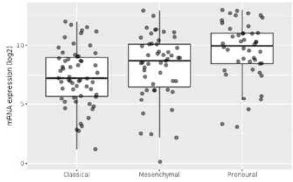


| | Sample_count | median | mad | mean | sd |
|---|---|---|---|---|---|
| Classical | 59 (37.82%) | 7.20 | 2.41 | 7.33 | 2.44 |
| Mesenchymal | 51 (32.69%) | 8.69 | 2.54 | 8.19 | 2.73 |
| Proneural | 46 (29.49%) | 9.96 | 2.16 | 9.51 | 2.49 |
| TOTAL | 156 | 8.62 | 2.66 | 8.25 | 2.69 |

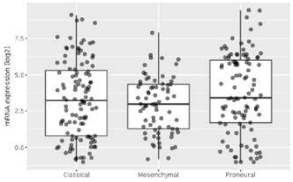


| | Sample_count | median | mad | mean | sd |
|---|---|---|---|---|---|
| Classical | 111 (38.68%) | 3.23 | 3.36 | 3.32 | 2.65 |
| Mesenchymal | 73 (25.44%) | 2.97 | 2.31 | 2.92 | 1.94 |
| Proneural | 103 (35.89%) | 3.44 | 3.50 | 3.71 | 2.74 |
| TOTAL | 287 | 3.33 | 3.04 | 3.36 | 2.53 |

**F** CD248

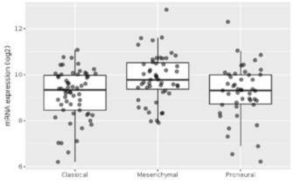


| | Sample_count | median | mad | mean | sd |
|---|---|---|---|---|---|
| Classical | 59 (37.82%) | 9.33 | 1.01 | 9.14 | 1.07 |
| Mesenchymal | 51 (32.69%) | 9.78 | 0.99 | 9.86 | 1.03 |
| Proneural | 46 (29.49%) | 9.31 | 0.96 | 9.23 | 1.16 |
| TOTAL | 156 | 9.50 | 0.94 | 9.40 | 1.13 |

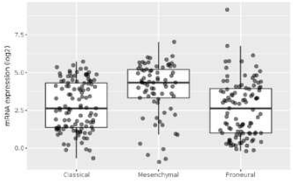


| | Sample_count | median | mad | mean | sd |
|---|---|---|---|---|---|
| Classical | 111 (38.68%) | 2.64 | 2.14 | 2.80 | 1.63 |
| Mesenchymal | 73 (25.44%) | 4.34 | 1.37 | 3.94 | 1.75 |
| Proneural | 103 (35.89%) | 2.64 | 2.27 | 2.59 | 1.86 |
| TOTAL | 287 | 3.29 | 2.26 | 3.02 | 1.82 |

**G** CD31

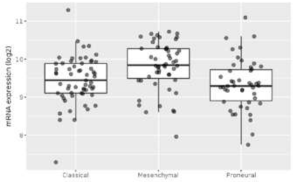


| | Sample_count | median | mad | mean | sd |
|---|---|---|---|---|---|
| Classical | 59 (37.82%) | 9.44 | 0.55 | 9.45 | 0.61 |
| Mesenchymal | 51 (32.69%) | 9.84 | 0.60 | 9.81 | 0.64 |
| Proneural | 46 (29.49%) | 9.30 | 0.63 | 9.34 | 0.66 |
| TOTAL | 156 | 9.59 | 0.60 | 9.53 | 0.66 |

No available data.

**Supplementary Figure S1. Subtype specific expression patterns from TCGA and CGGA databases.**
(A-G) RNAseq expression patterns seen in classical, proneural, or mesenchymal type GBM. Results reflect GSC health (A-C) and differentiation markers (D-G).

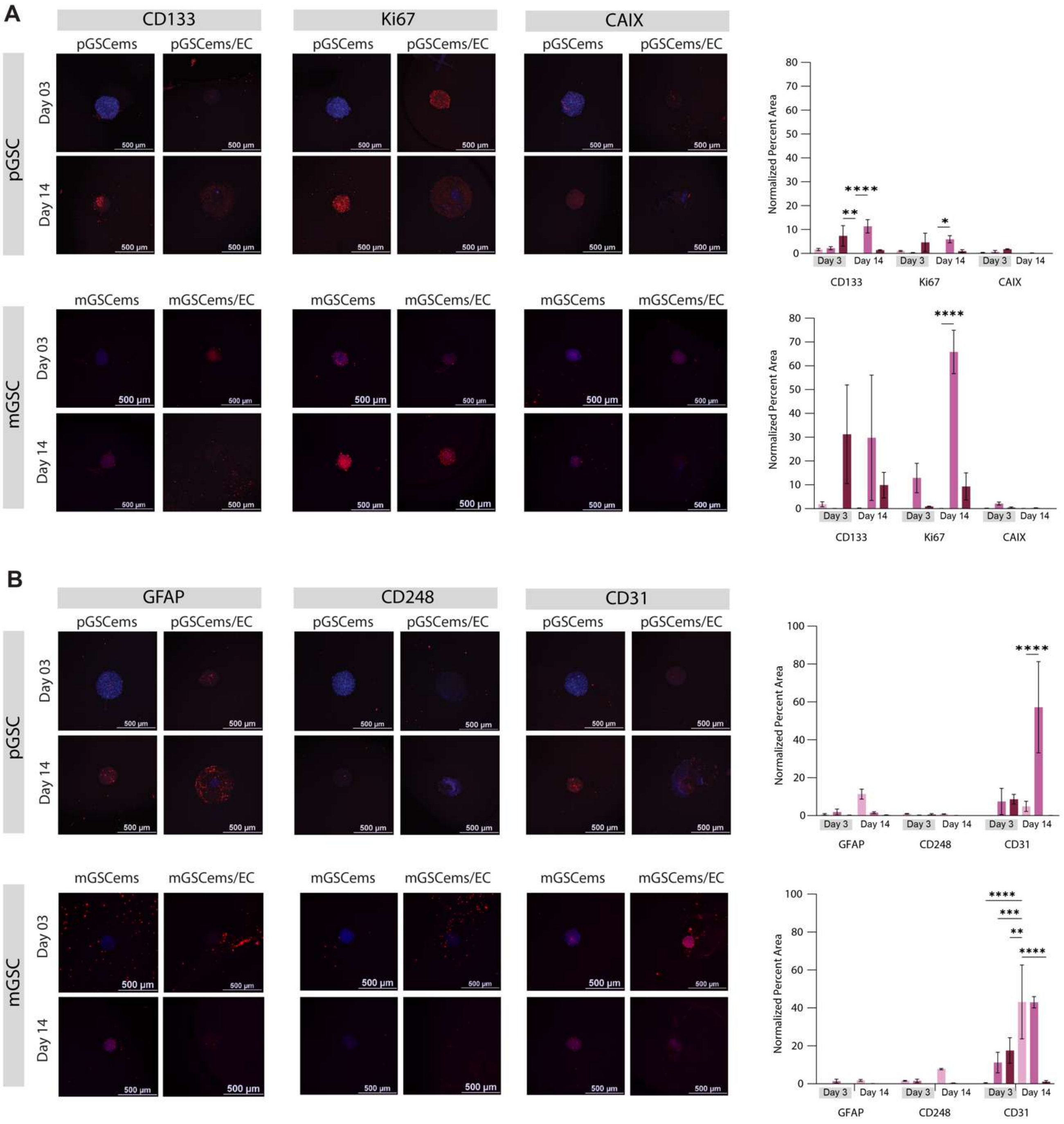


**Supplementary Figure S2. Expanded IF staining results seen in Figure 6.** (A) Immunofluorescence (IF) staining and quantification of stemness (CD133), proliferation (Ki67), and hypoxia (CAIX) markers in pGSC and mGSC spheroids under encapsulated and co-culture conditions. (B) IF staining and quantification of astrocytic (GFAP), oligodendrocytic (MOG), pericytic (CD248), and endothelial (CD31) differentiation markers, highlighting enhanced lineage specification with EC co-culture. Data are presented as mean ± SD (N = 3). Statistical analysis was performed using two-way ANOVA with Tukey's post hoc test (*p ≤ 0.05, **p ≤ 0.005, ***p ≤ 0.0005, ****p ≤ 0.00005).
